# The activated plant NRC4 immune receptor forms a hexameric resistosome

**DOI:** 10.1101/2023.12.18.571367

**Authors:** Furong Liu, Zhenlin Yang, Chao Wang, Raoul Martin, Wenjie Qiao, Jan E. Carette, Sheng Luan, Eva Nogales, Brian Staskawicz

## Abstract

Innate immune responses against microbial pathogens in both plants and animals are regulated by intracellular receptors known as Nucleotide-binding Leucine-rich Repeats (NLR) proteins. In plants, these NLRs play a crucial role in recognizing pathogen effectors, thereby initiating the activation of immune defense mechanisms. Notably, certain NLRs serve as “helper” NLR immune receptors (hNLR), working in tandem with “sensor” NLR immune receptors (sNLR) counterparts to orchestrate downstream signaling events to express disease resistance. In this study, we reconstituted and determined the cryo-EM structure of the hNLR required for cell death 4 (NRC4) resistosome. The auto-active NRC4 formed a previously unanticipated hexameric configuration, triggering immune responses associated with Ca^2+^ influx into the cytosol. Furthermore, we uncovered a dodecameric state of NRC4, where the coil-coil (CC) domain is embedded within the complex, suggesting an inactive state, and expanding our understanding of the regulation of plant immune responses.

**One Sentence Summary:** The hexameric NRC4 resistosome mediates cell death associated with cytosolic Ca^2+^ influx.

## Main Text

Plants and animals rely on innate immune receptors to detect and respond to invading pathogens. In plants, these receptors are primarily of two types: cell-surface pattern recognition receptors (PRRs) and intracellular NLRs (*1, 2*). PRRs recognize microbe/pathogen-associated molecular patterns and activate pattern-triggered immunity (PTI), while NLRs detect pathogen effectors either directly or indirectly by recognizing alterations of host targets, ultimately leading to the activation of effector-triggered immunity (ETI) (*1*). Recent research has unveiled an intricate interplay between ETI and PTI in plants (*3-5*). It has been revealed that NLRs not only execute their direct roles in ETI but also significantly contribute to the enhancement of PTI by upregulating PTI signaling components (*3-5*).

Pathogen effector recognition leads to the oligomerization of the NLRs acting as sensors for effectors that directly bind immune receptor (*6-10*). For example, recognition of the pathogen effectors XopQ1 and ATR1 by TIR domain-containing NLR (TNL)-type resistance proteins ROQ1 and RPP1 leads to the formation of tetrameric resistosomes respectively (*7, 8*). These tetramers activate nicotinamide adenine dinucleotide nucleosidase (NADase) activity of the TIR domain, generating small molecules with the potential to serve as second messengers, activating downstream immune responses (*11-13*). On the other hand, Arabidopsis HOPZ-ACTIVATED REASISTANCE 1 (ZAR1) and wheat stem rust resistance 35 (Sr35) have been shown to form pentameric resistosomes upon effector detection (*6, 9, 10*). Within these two resistosomes, the CC domains insert into the plasma membrane (PM), forming calcium (Ca^2+^) permeable cation channels, resulting in robust Ca^2+^ influx across the PM and elevation in cytosolic Ca^2+^ ([Ca^2+^]_cyt_), essential for inducing defense responses and cell death (*9, 14*).

Specific NLRs, referred to as ‘helper’ NLRs (hNLRs), are essential for collaborating with and facilitating certain sensor NLRs (sNLRs) to regulate both PTI and ETI responses (*15-19*). NRCs are well-know hNLRs in *Solanaceous* plants with capability to transmit immune signals from various sNLRs and specific PRRs (*18, 20*). Upon the effector recognition by sNLRs, NRC0, NRC2, NRC3 and NRC4 have been shown to form oligomerized macromolecules (*21-23*). Subsequently, these oligomers initiate ETI, resulting in a programmed cell death known as hypersensitive response (HR) that is associated with the limitation of pathogen proliferation (*2*). In addition, to autoactivate hNLRs, N REQUIRED GENE1 (NRG1) and ACTIVATED DISEASE RESISTANCE1 (ADR1) oligomerize and form Ca^2+^-permeable cation channels, leading to an increase in [Ca^2+^]_cyt_ levels and cell death (*24*). However, it is still unclear whether NRCs’ activation share the same function related to Ca^2+^ influx.

Recently, multiple cryo-EM studies have provided valuable insights into the structures of the sNLR resistosomes (*6-10*). However, the structures of the hNLRs are still unknown, limiting our understanding of the molecular basis, activation, and regulation of these indispensable components in plant immune responses.

## Results

### Reconstitution and cryo-electron microscopy structure of the NRC4 resistosome

NRC4 follows the typical CNL domain structure, comprising an N-terminal CC domain, followed by a Nucleotide-Binding domain (NBD), Helical Domain 1 (HD1), a Winged-Helix Domain (WHD), and a C-terminal Leucine-Rich-Repeat (LRR) domain (Fig.1A) (*25*). We isolated the NRC4 resistosome from *Nicotiana benthamiana* (*N. benthamiana*) leaves through the introduction of two specific mutations. Mutation D478V (DV) has been shown to be an autoactive mutation, while mutation L9E inhibits the HR cell death induced by NRC4 DV (Fig.S1) (*25*), thus ensuring the formation of an activated NRC4 resistosome without causing cell death. Inoculation of *N. benthamiana* leaves with two vectors carrying either a 3X Flag or a Twin-StrepII tag, in conjunction with the double-mutated NRC4 protein, facilitated the successful tandem affinity purification of highly pure NRC4 protein (Fig.S2). This purification strategy proved exceptionally efficient, suggesting an oligomerization state of NRC4 that was further confirmed through structural analysis. Cryo-EM samples of NRC4 were prepared on grids with either carbon or graphene support layers as a means to concentrate the resistosome for visualization. Early image analysis showed two distinct views, corresponding to either one or two density layers (Fig.S3A). The double-layer architecture represents a significant departure from the prevailing resistosome structural model, typically featured as single-layered (*6-10*). Further analysis resulted in two distinct NRC4 resistosome structures obtained from the same sample: a 2.6 Å hexameric structure (Fig.1B and C), and a 3.5 Å dodecameric structure (Fig.1D).

**Fig 1.**
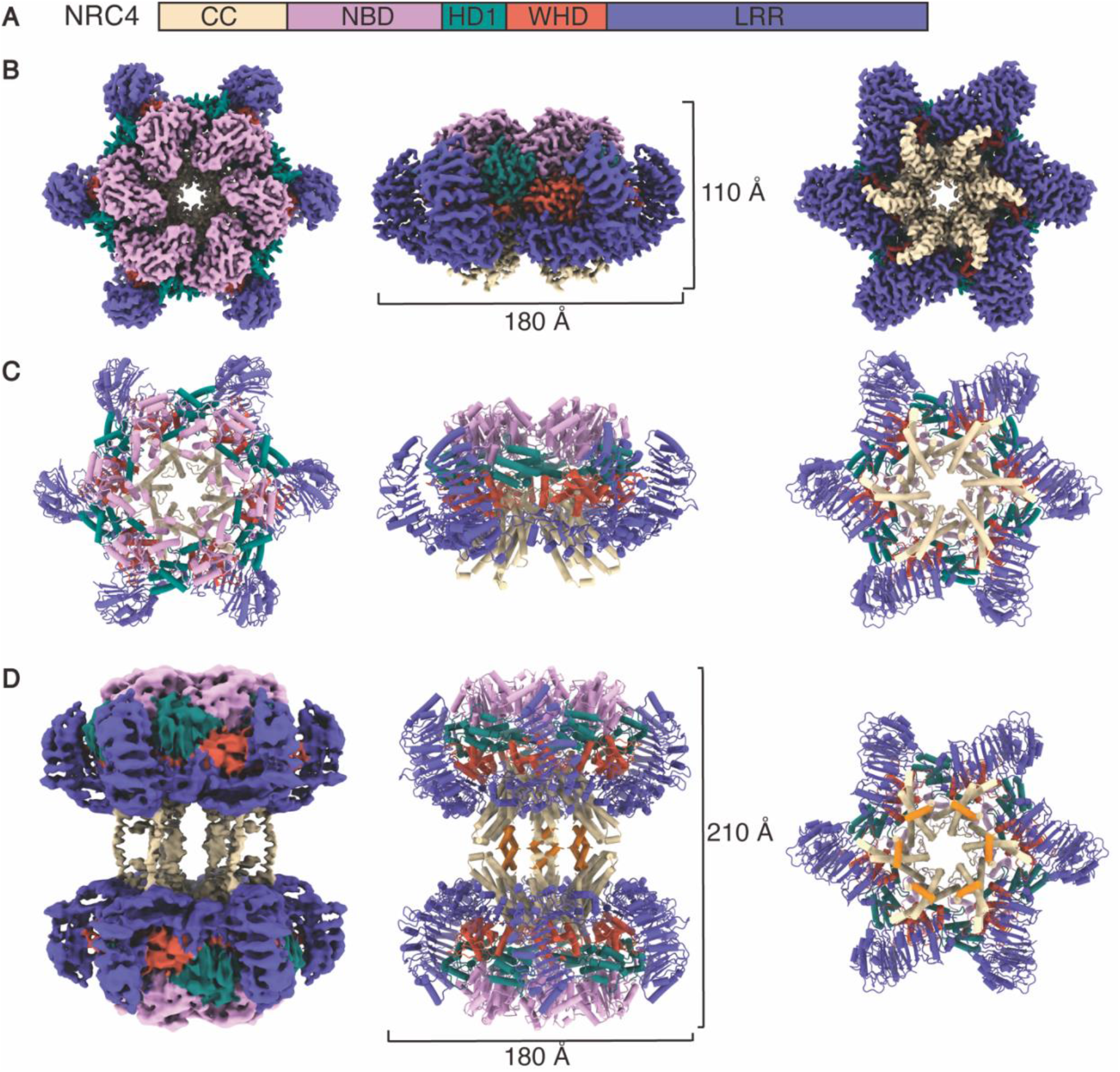
Structure of the NRC4 resistosome. **(A)** Domain organization of NRC4. The same color code for the domains is used throughout this study, unless otherwise specified. **(B)** Cryo-EM density map of the hexameric NRC4 structure, and **(C)** corresponding refined structure model (top, side and bottom views are shown from left to right). Six identical NRC4 protomers are arranged in a wheel-like structure measuring ∼180 Å in diameter and 100 Å in height. **(D)** Left, cryo-EM density map of the dodecameric NRC4 resistosome; middle, corresponding atomic model; right, cross-sectional view of the dodecameric structural model. Twelve identical NRC4 protomers form a dumbbell-shaped structure, measuring ∼180 Å in diameter and 210 Å in height. The α1-helixes in the CC domains, which bridge the formation of the double-layer structure, are highlighted in orange.

### Overall structure of NRC4 resistosome

The resolution of the cryo-EM maps allowed us *de novo* atomic models building of the NRC4 oligomers, which exhibit architectural features akin to well-established CNLs structures that are pentamers (*6, 9, 10*). The hexameric assembly of NRC4 results in a wheel-like structure measuring approximately 180 Å in diameter and 100 Å in height (Fig.1B). Compared to that of the ZAR1 and Sr35 resistosomes, the organization of the NRC4 hexamer involves a more densely packed arrangement of protomers (Fig.1B and C). This dense packing, particularly of the LRR domain, limits the available space for interactions with external protein factors, as seen for effectors in the case of ZAR1 and Sr35 (*6, 9*). The LRR domain in NRC4 arranges at a smaller angle relative to the tangent of the wheel-like structure (Fig.1B and C), allowing the LRR to extend more prominently into the solvent, and thus compensating for the restricted space between adjacent LRRs (Fig.1B and C). In the hexameric NRC4 resistosome, the α_1_ helix in the CC domain, equivalent to that seen for Sr35 (*9*), was not visible, likely due to flexibility.

In addition to the single-layer wheel-like structure previously seen for other plant resistosomes, we identified an alternative state of the NRC4 resistosome in which two NRC4 hexamers face each other forming a dumbbell-shaped dodecamer (Fig.1D and Fig.S3), measuring approximately 180 Å in diameter and 210 Å in height. The connections between the two hexamers were assigned to two antiparallel α-helices corresponding to the α1-helix from the CC domain as predicted by AlphaFold multimer that we could unambiguously fit into the cryo-EM map (Fig.1D and Fig.S4) (*26, 27*). Thus, our analysis underscores the coexistence of two distinct NRC4 resistosome states, a classical single-layer wheel-like structure (Fig.1B and C), and a non-classical double-layer configuration (Fig.1D) in which the layers are bridged by the N-terminus α_1_-helices of the protomers.

### Oligomerization of the NRC4 resistosome

Protomer packing in the NRC4 resistosome structure is governed by multiple interactions involving the five domains (Fig.2A). In the interaction between adjacent CC domains, the α2-helix from one protomer engages with the α3-helix of its neighboring protomer (Fig.2B). D47 from α2-helix hydrogen bonds with mainchain atoms of both E56 and E57 in the α3-helix (Fig.2B) and Q129 from the α2-helix hydrogen bonds with the main chain of Q126 in the α3-helix (Fig.2B). Interestingly, the NBD-NBD interaction, previously implicated in protomer packing and contributing to Sr35 and ZAR1 resistosome formation (*6, 9*), was not detected in the NRC4 structure. Instead, we observed that amino acids E283 and D275 from the NBD domain form two salt bridge interactions with K329 and R262, respectively, in the HD1 domain of the neighboring protomer (Fig.2C). D453 from the WHD is also engaged in a salt bridge interaction with K364 from the HD1 domain (Fig.2D). Similar to ZAR1, but unlike Sr35, the LRR domains pack directly against each other, with a hydrogen bond between C661 and D502 (Fig.2E) (*6, 9*). To assess the significance of these interactions in facilitating the NRC4-mediated HR cell death, we introduced single point mutations and group mutations in these interfaces. Two groups, H40A/D47A/Q129A and D275A/E283A/D453A, completely abolished cell death activity (Fig.2G). Of note, the single mutation D47A was capable of abolishing cell death on the *N. benthamiana* leaves, highlighting the importance of this residue in NRC4’s function (Fig.2G). Both wild-type NRC4 and all tested mutants that hinder HR exhibited comparable protein levels, indicating that these substitutions do not affect NRC4 stability (Fig.2H).

**Fig 2.**
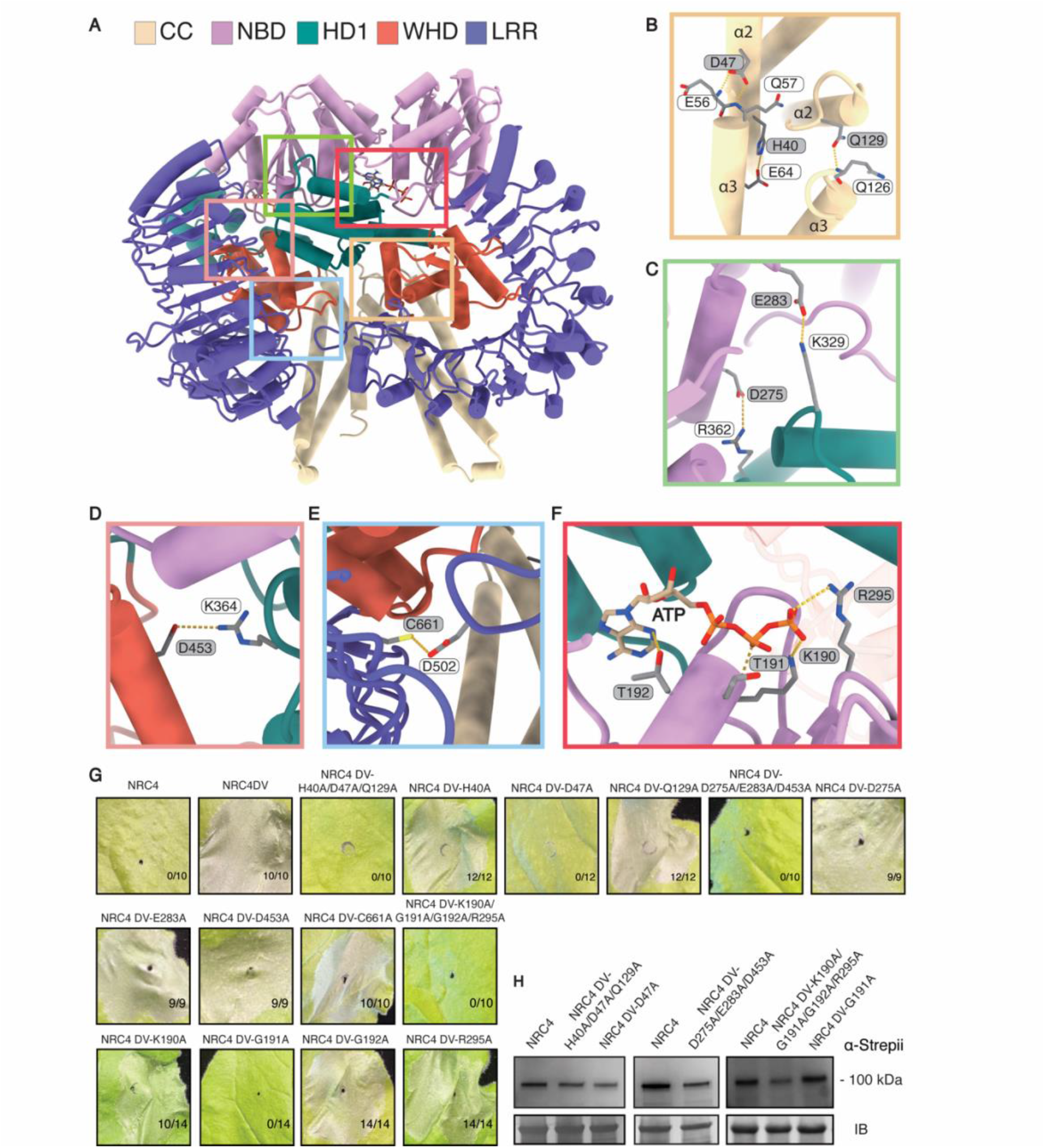
Interfaces in the oligomerization of the NRC4 resistosome. **(A)** View of two adjacent protomers in the NRC4 resistosome. Boxes indicate regions of interaction between them shown in detail of panels B-E. **(B-E)** Structural details of CC-CC, NBD-HD, WHD-HD1 and LRR-LRR interactions, respectively. Residue labels in both grey and white correspond to two adjacent protomers. **(F)** ATP is situated in the cleft between the NBD and HD1 domain. Although it is in proximity to both domains, ATP interacts exclusively with the NBD. **(G)** Hypersensitive response phenotypes of *N. benthamiana* leaves upon the expression of mutant NRC4 created based on the interfaces shown in (B-F). In each case a representative figure is shown from multiple replicates. **(H)** The protein expression levels of the wild type NRC4 and the tested mutants in the *N. benthamiana* leaves were evaluated using SDS-PAGE and subsequent immunoblotting with α-Strepii antibody. InstantBlue® Coomassie-stained gel was used as loading control (IB, InstantBlue).

ATP binding leads to a sequence of structural rearrangements crucial for NLRs to switch from inactive to active states (*6, 9*). Our NRC4 structure shows clear density for an ATP molecule located within the cleft formed by the NBD and HD1 domains (Fig.2F). This position resembles that seen for Sr35 but differs from the configuration in the ZAR1 resistosome, where ATP predominantly interacts with the WHD domain (*6, 9, 10*). In NRC4, ATP only interacts with residues in the NBD, despite its proximity to the HD1 domain (Fig.2F). Specifically, T192 hydrogen bonds with the adenine base of ATP, while K190, T191, and R295 interact with the ATP phosphates (Fig.2F). Mutations disrupting these interactions (K190A/T191A/T192A/R295A) led to the inhibition of HR in *N. benthamiana* leaves (Fig.2G). A significant loss of HR was also observed with the single point mutation T191A (Fig.2G), emphasizing its essential role in ATP binding. Notably, the protein levels of wild-type NRC4 and all tested mutants that hinder HR remained comparable, indicating that these substitutions do not affect the stability of NRC4 (Fig 2H).

Previous cryo-EM studies have elucidated high-resolution single-layer plant resistosome structures (*6-10*). However, in the study of the Sr35 resistosome, 2D analysis showed the presence also of a double-layer structure where the two layers appear flexibly attached (*9, 10*). We were able to obtain the double-layer structure of the NRC4 resistosome in our study (Fig.1D and Fig.S3), clearly revealing a higher oligomerization state of the plant resistosome. Surprisingly, we found the CC domains, which are crucial for cell death, locked inside the dodecameric structure. While in the hexameric structure the CC domains are exposed (Fig.1B and 1C) and could therefore interact with the plasma membrane, in the dodecameric configuration interaction of the two CC domains on opposite sides of the double-layer with each other would prevent the NRC4 resistosome from anchoring to the plasma membrane. In this configuration the hydrophobic residues in the α1-helices of the CC domains that would insert in the lipid bilayer are interacting with each other as they bring the two hexamers together (Fig.S4A)(*26, 27*). This interaction was confirmed by a co-immunoprecipitation (co-IP) assay (Fig.S4B), where the flag-tagged α1-helix was able to pull down the Strep-tagged α1-helix, *vice versa*. Thus, we attribute the higher-order oligomerization state of the dodecameric NRC4 resistosome to an inactive formation.

### Conservation of the EDVID motif and LRR^R-cluster^ in NRC4

The EDVID sequence motif in the α3-helix is engaged with a positively charged region on the LRR domains known as the R-cluster motif, resulting in an intramolecular interactions that have been shown to be critical for CNLs-mediated cell death (*9, 28*). In the NRC4 resistosome structure, only this α3-helix interacts with the LRR domain, while the other helices within the CC domain do not participate in such interaction (Fig.3A). Residues E73, D74, and D77 within the α3-helix form salt bridges and hydrogen bonds with R514 and R537 of the LRR domain (Fig.3B). An electrostatic surface analysis of the LRR domain in NRC4 revealed a striking similarity in size to the R-cluster observed in ZAR1 and Sr35 (Fig.3C). Sequence alignment analysis further revealed the conservation of the EDVID motif when comparing NRCs with ZAR1 and Sr35 (Fig.3D). The two arginine residues (R514 and R537) in the R-cluster of NRC4 involved in the interaction with CC domain are also present in Sr35 (Fig.3D). While in both of the ZAR1 and Sr35 resistosomes structures five arginine residues from distinct repeats within the LRR domain engage in extensive interactions with the CC domain (*6, 9*), fewer arginines in the LRR domain of NRC4 interact with the CC domain (Fig.3B and D). To assess the importance of the EDVID motif and the R-cluster in the immune response, we mutated key amino acids participating in this interaction. We found both triple mutations of E73A/D74A/D77A in the EDVID motif and double mutations of R514A/R537A in the R-cluster resulted in a loss of cell death activity (Fig.3E), without affecting the stability of NRC4 protein (Fig.3F). Single amino acid substitutions of E73, D74, or D77 failed to abolish cell death activity (Fig.3E), while the single R514A or R537A substitution was enough to hinder cell death activity (Fig.3F), emphasizing the crucial role of these two arginines within the R-cluster (Fig. 3B and D). These findings suggest an essential involvement of the EDVID and R-cluster motifs in NRC4 DV-mediated cell death.

**Fig 3.**
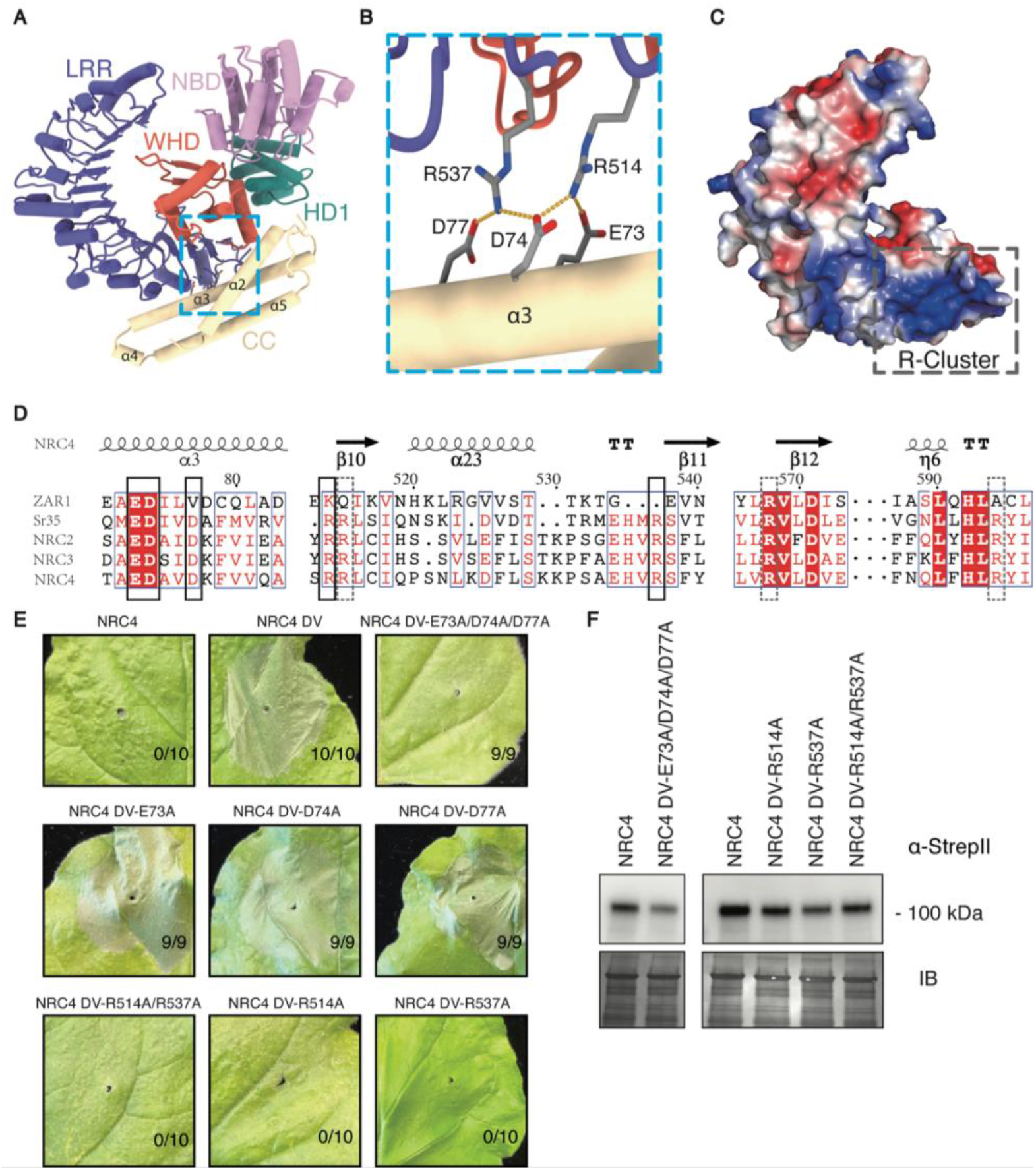
CC-LRR interactions within an NRC4 protomer. **(A)** Structure of one NRC4 protomer, with the CC-LRR interaction region indicated by the light blue dashed box. **(B)** Structural details of the CC-LRR interactions. Three negatively charged residues (E73, D74, and D77) from the CC domain form salt bridge and hydrogen bond interactions with two positively charged residues (R514 and R537) from the LRR domain. **(C)** Electrostatic surface view of the LRR domain. The position of the conserved R-cluster is indicated by the dashed box. **(D)** Structure-based sequence alignment encompassing the EDVID motif and R-cluster of Zar1, Sr35, NRC2, NRC3 and NRC4. Key residues shown in (B) are highlighted by a solid box, while the additional arginines involved in the CC-LRR interaction of the Sr35 resistosome are highlighted by a dashed box. **(E)** Hypersensitive response phenotypes of *N. benthamiana* leaves upon the expression of NRC4 with mutations of the residues shown in (B). In each case, one representative figure is shown from multiple replicates. **(F)** The protein expression levels of the wild type NCR4 and the tested mutants in the *N. benthamiana* leaves were evaluated using SDS-PAGE and subsequent immunoblotting with α-Strepii antibody. InstantBlue® Coomassie-stained gel was used as loading control.

### Autoactive NRC4 triggers calcium influx

Increasing evidence underscores the critical role of Ca^2+^ signaling in the initiation of ETI (*29*). Particularly relevant are the findings that both CNLs (e.g., ZAR1 and Sr35) and hNLRs (e.g., NRG1.1) assemble into pore-forming resistosomes within the PM, facilitating the influx of extracellular Ca^2+^ into the cytosol (*9, 14, 24, 30, 31*). We thus hypothesized that the activation of NRC4 may trigger a Ca^2+^ influx that appears to be a common downstream event of NLR activation. To test this hypothesis, we transiently expressed several NRC4 variants, including the autoactive NRC4 DV, in *N. benthamiana* leaves expressing the [Ca^2+^]_cyt_ reporter GCaMP3 (*18, 32*). Our results revealed a transient [Ca^2+^]_cyt_ increase approximately 8-9 hours after leaf infiltration with Agrobacterium carrying vectors expressing the active NRC4 DV (Fig 4A and 4B). Notably, this [Ca^2+^]_cyt_ increase preceded the onset of leaf HR cell death, which typically occurred over a more extended time frame (Fig.S4). In contrast, the wild-type NRC4, or the variants NRC4 L9E and NRC4 DV/L9E, failed to induce either [Ca^2+^]_cyt_ increase (Fig 4A and 4B), or cell death (Fig.S1A). Furthermore, application of a Ca^2+^-channel blocker, Lanthanum (III) chloride (LaCl_3_), effectively abolished NRC4 DV-mediated [Ca^2+^] influx (Fig.4C and 4D) and HR cell death (Fig.S5A). In contrast, elevated external Ca^2+^ enhanced NRC4 DV-mediated Ca^2+^ influx (Fig.4E and 4F). To explore this possible mechanism, we attempted to assess the channel function of NRC4 DV in a mammalian HEK239FT cell line. Unexpectedly, and in contrast to NRG1.1 DV, NRC4 DV failed to associate with the PM (Fig.S6A). In addition, NRG1.1 DV, but not NRC4 DV, induced growth defects and substantial cell death in this mammalian cell (Fig.S6B) (*24*). Together, these results support the conclusion that the active NRC4 resistosome facilitates the influx of extracellular Ca^2+^ into the cytosol of plant cells, a pivotal prerequisite for NRC4 resistosome-mediated HR cell death.

**Fig 4.**
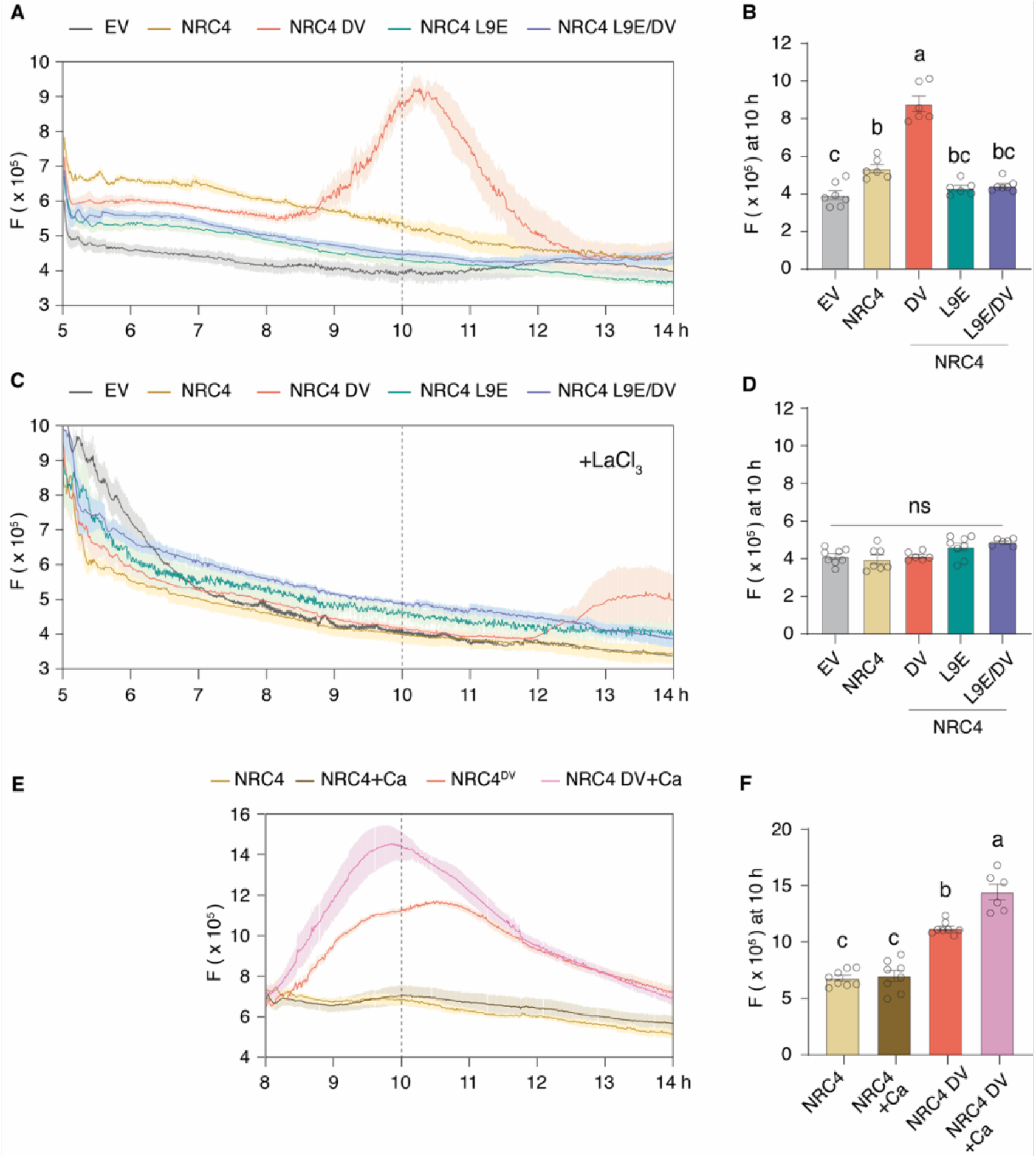
[Ca^2+^]_cyt_ dynamics upon the expression of NRC4 variants and indicated treatments. **(A)** Time course of [Ca^2+^]_cyt_ dynamics after infiltration of *N. benthamiana* leaves expressing the [Ca^2+^]_cyt_ reporter GCaMP3 with Agrobacterium strains carrying the indicated constructs. NRC4 DV and NRC4 L9E denote NRC4 variants with amino acid the substitutions D478V and L9E, respectively. NRC4 L9E/DV indicates an NRC4 variant with both mutations. Leaf disc fluorescent (F) intensities of GCaMP3, as indicative of relative [Ca^2+^]_cyt_ levels, are plotted over a tested time. The autoactive NRC4 DV, unlike other variants, exhibited a robust increase in [Ca^2+^]_cyt_. **(B)** Relative [Ca^2+^]_cyt_ levels at indicated time as shown in (A). **(C)** Time course of [Ca^2+^]_cyt_ dynamics upon the expression of NRC4 variants with co-infiltration with a PM calcium channel block, LaCl_3_ (2nM). **(D)** Relative [Ca^2+^]_cyt_ levels at indicated time as shown in (C). **(E)** Time course of [Ca^2+^]_cyt_ dynamics, showing additive effects of extracellular Ca^2+^ (10 mM) on NRC4 DV-mediated Ca^2+^ influx. (F) Relative [Ca^2+^]_cyt_ levels at indicated time as shown in (E). Error bars in panels A to F represent standard error (n=6 to 8 discs from 3 independent plants). Experiments were repeated twice with similar phenotypes observed.

## Discussion

### Distinct activation mechanisms of sensor-helper NLR pairs in mammals and plants

In mammalian cells, the NLR family of apoptosis inhibitory proteins (NAIPs) act as sNLRs and are activated by binding to specific bacterial protein ligands (*33*). Upon activation, NAIPs co-assemble with a downstream hNLR, the NLR family CARD domain-containing protein 4 (NLRC4), forming NAIP-NLRC4 inflammasomes, which play crucial roles in the innate immune system (*34-36*). Each inflammasome compromise one NAIP and multiple NLRC4 protomers (*33, 37*). Our study shows that NRC4 oligomerizes without the need to incorporate sNLRs. The presence of six identical protomers in the NRC4 resistosome supports an activation-and-release model for plant sNLR/hNLR signaling, wherein hNLRs do not form stable complex with sNLRs (*21, 22*). Thus, our findings highlight the significant differences in the activation mechanisms of sNLR/hNLR pairs between plants and animals.

### Dual-hexamer formation and immune regulation

A feature clearly unveiled in our study of the NRC4 resistosomes was the formation of both single-layer, hexameric assemblies, and double-layer, dodecameric assemblies. In our dodecameric NRC4 resistosome structure, the CC domain, which is crucial for triggering cell death, is locked at the center of the dodecamer (Fig.1D). We noticed that in the prior studies of the Sr35 resistosome, 2D classification also showed the presence of double-layer states alongside the single-layer configuration (*9, 10*). Therefore, an inactive, higher-order oligomer might be a common feature in plant resistosomes, providing a mechanism for the regulation of plant immune responses. This intriguing phenomenon is reminiscent of mouse NLRC4, which was also found to form a double-layer assembly in which its caspase activation and recruitment domains (CARDs) was buried at the center (*33*), potentially restraining its role in immune response. Interestingly, single point mutations in the WHD domain of NLRC4 identified as gain-of-function mutations that result in auto-inflammatory diseases in humans are positioned in a similar location as the DV mutation in the NRC4 WHD domain that triggers cell death (Fig.S1) (*25, 38-41*). Given the structural and functional similarities observed in the double-layer configurations, we hypothesize that the dodecameric state of the NRC4 resistosome may serve as a “protector” against undesired immune activation provoked by harmful mutations. Alternatively, this state may function as a “reservoir” for storing NRC4 protein without triggering an immune response.

### Plant NLR signaling: insights into Ca^2+^ influx as a central mechanism

Both CNL- and TNL-mediated immune responses in plant cells converge on Ca^2+^ influx, indicating that Ca^2+^ serves as a common messenger for plant NLR signaling. Previous studies have shown that two sNLR resistosomes, ZAR1 and Sr35, as well as the hNLR NRG1 oligomer, can form Ca^2+^ permeable channels (*9, 14, 24*). Duggan *et al*. reported that the NRC4 DV resistosome exhibits a punctate distribution that is primarily associated with the PM, a pattern similar to that of the hNLR NRG1 resistosome (*42*). Indeed, NRC4 DV, like NRG1.1 DV, triggered a robust Ca^2+^ influx when expressed in plants. Taken together, these findings suggest that the NRC4 DV resistosomes may also form a Ca^2+^-permeable channel in plant, thereby triggering immune responses and cell death. But NRC4 DV functions a different way in human cells as NRG1.1 DV based on our preliminary data, then it is possible that NRC4 DV requires other unidentified plant-specific regulators (absent in mammalian cells) for its effective association with the PM. This might represent a unique activation mechanism of NRC hNLRs distinct from that of the NRG1 helper NLRs and warrants future research.

In summary, we describe an activated hexameric NRC4 resistosome, potentially forming a Ca^2+^ permeable channel to trigger immune response. We also reveal a dodecameric assembly of the NRC4 resistosome with the CC domain locked away within the structure, presumably preventing its anchoring to the PM. Our combined data provide a mechanistic model of hNLR resistosome formation and regulation, offering the promise of advancing our basic understanding of plant pathogen defense.

## Acknowledgments

We thank R. Vance for his helpful comments and suggestions, D. Toso, R. Thakkar and P. Tobias at the Cal-Cryo facility for their assistance with cryo-EM data collection and handling, K. Stine for computational support, and P. Cao for advice on cryo-EM grid preparation and data analysis. Transgenic *N. benthamiana* seeds carrying GCaMP3 were kindly provided by K. Yoshioka.

## Author contributions

Conceptualization: F.L., Z.Y., C.W., B.J.S.; Methodology: F.L., Z.Y., C.W., R.M.; Investigation: F.L., Z.Y., C.W., W.Q; Visualization: F.L., Z.Y., C.W., W.Q; Funding acquisition: B.J.S., S.L., E.N.; Supervision: B.J.S., E.N.; Writing – original draft: F.L., Z.Y., C.W.; Writing – review & editing: F.L., Z.Y., C.W., J.E.C., S.L., E.N., B.J.S.

## Data and materials availability

Materials are available from B.J.S.

## References and notes

1. B. P. M. Ngou, P. Ding, J. D. Jones, Thirty years of resistance: Zig-zag through the plant immune system. The Plant cell 34, 1447–1478 (2022).

2. J. D. Jones, R. E. Vance, J. L. Dangl, Intracellular innate immune surveillance devices in plants and animals. Science 354, aaf6395 (2016).

3. M. Yuan et al., Pattern-recognition receptors are required for NLR-mediated plant immunity. Nature 592, 105–109 (2021).

4. B. P. M. Ngou, H.-K. Ahn, P. Ding, J. D. Jones, Mutual potentiation of plant immunity by cell-surface and intracellular receptors. Nature 592, 110–115 (2021).

5. H. Tian et al., Activation of TIR signalling boosts pattern-triggered immunity. Nature 598, 500–503 (2021).

6. J. Wang et al., Reconstitution and structure of a plant NLR resistosome conferring immunity. Science 364, eaav5870 (2019).

7. R. Martin et al., Structure of the activated ROQ1 resistosome directly recognizing the pathogen effector XopQ. Science 370, eabd9993 (2020).

8. S. Ma et al., Direct pathogen-induced assembly of an NLR immune receptor complex to form a holoenzyme. Science 370, eabe3069 (2020).

9. A. Förderer et al., A wheat resistosome defines common principles of immune receptor channels. Nature 610, 532–539 (2022).

10. Y.-B. Zhao et al., Pathogen effector AvrSr35 triggers Sr35 resistosome assembly via a direct recognition mechanism. Science Advances 8, eabq5108 (2022).

11. S. Huang et al., Identification and receptor mechanism of TIR-catalyzed small molecules in plant immunity. Science 377, eabq3297 (2022).

12. A. Jia et al., TIR-catalyzed ADP-ribosylation reactions produce signaling molecules for plant immunity. Science 377, eabq8180 (2022).

13. D. Yu et al., TIR domains of plant immune receptors are 2′, 3′-cAMP/cGMP synthetases mediating cell death. Cell 185, 2370–2386. e2318 (2022).

14. G. Bi et al., The ZAR1 resistosome is a calcium-permeable channel triggering plant immune signaling. Cell 184, 3528–3541. e3512 (2021).

15. V. Bonardi et al., Expanded functions for a family of plant intracellular immune receptors beyond specific recognition of pathogen effectors. Proceedings of the National Academy of Sciences 108, 16463–16468 (2011).

16. J. R. Peart, P. Mestre, R. Lu, I. Malcuit, D. C. Baulcombe, NRG1, a CC-NB-LRR protein, together with N, a TIR-NB-LRR protein, mediates resistance against tobacco mosaic virus. Current biology 15, 968–973 (2005).

17. J. M. Feehan et al., Oligomerization of a plant helper NLR requires cell-surface and intracellular immune receptor activation. Proceedings of the National Academy of Sciences 120, e2210406120 (2023).

18. J. Kourelis et al., The helper NLR immune protein NRC3 mediates the hypersensitive cell death caused by the cell-surface receptor Cf-4. PLoS genetics 18, e1010414 (2022).

19. R. N. Pruitt et al., The EDS1–PAD4–ADR1 node mediates Arabidopsis pattern-triggered immunity. Nature 598, 495–499 (2021).

20. C.-H. Wu et al., NLR network mediates immunity to diverse plant pathogens. Proceedings of the National Academy of Sciences 114, 8113–8118 (2017).

21. M. P. Contreras et al., Sensor NLR immune proteins activate oligomerization of their NRC helpers in response to plant pathogens. The EMBO Journal 42, e111519 (2023).

22. H. K. Ahn et al., Effector-dependent activation and oligomerization of plant NRC class helper NLRs by sensor NLR immune receptors Rpi-amr3 and Rpi-amr1. The EMBO Journal 42, e111484 (2023).

23. T. Sakai et al., The NRC0 gene cluster of sensor and helper NLR immune receptors is functionally conserved across asterid plants. bioRxiv, x2023.2010. 2023.563533 (2023).

24. P. Jacob et al., Plant “helper” immune receptors are Ca2+-permeable nonselective cation channels. Science 373, 420–425 (2021).

25. H. Adachi et al., An N-terminal motif in NLR immune receptors is functionally conserved across distantly related plant species. Elife 8, e49956 (2019).

26. J. Jumper et al., Highly accurate protein structure prediction with AlphaFold. Nature 596, 583–589 (2021).

27. M. Varadi et al., AlphaFold Protein Structure Database: massively expanding the structural coverage of protein-sequence space with high-accuracy models. Nucleic acids research 50, D439–D444 (2022).

28. G. J. Rairdan et al., The coiled-coil and nucleotide binding domains of the potato Rx disease resistance protein function in pathogen recognition and signaling. The Plant cell 20, 739–751 (2008).

29. C. Wang, S. Luan, Calcium homeostasis and signaling in plant immunity. Current opinion in plant biology 77, 102485 (2024).

30. M. Grant et al., The RPM1 plant disease resistance gene facilitates a rapid and sustained increase in cytosolic calcium that is necessary for the oxidative burst and hypersensitive cell death. The Plant Journal 23, 441–450 (2000).

31. X. Gao et al., Bifurcation of Arabidopsis NLR immune signaling via Ca2+-dependent protein kinases. PLoS pathogens 9, e1003127 (2013).

32. T. A. DeFalco et al., Using GCaMP3 to study Ca2+ signaling in Nicotiana species. Plant and Cell Physiology 58, 1173–1184 (2017).

33. J. L. Tenthorey et al., The structural basis of flagellin detection by NAIP5: A strategy to limit pathogen immune evasion. Science 358, 888–893 (2017).

34. E. F. Halff et al., Formation and structure of a NAIP5-NLRC4 inflammasome induced by direct interactions with conserved N-and C-terminal regions of flagellin. Journal of Biological Chemistry 287, 38460–38472 (2012).

35. L. Zhang et al., Cryo-EM structure of the activated NAIP2-NLRC4 inflammasome reveals nucleated polymerization. Science 350, 404–409 (2015).

36. Z. Hu et al., Structural and biochemical basis for induced self-propagation of NLRC4. Science 350, 399–404 (2015).

37. X. Yang et al., Structural basis for specific flagellin recognition by the NLR protein NAIP5. Cell research 28, 35–47 (2018).

38. S. W. Canna et al., An activating NLRC4 inflammasome mutation causes autoinflammation with recurrent macrophage activation syndrome. Nature genetics 46, 1140–1146 (2014).

39. A. Kitamura, Y. Sasaki, T. Abe, H. Kano, K. Yasutomo, An inherited mutation in NLRC4 causes autoinflammation in human and mice. Journal of Experimental Medicine 211, 2385–2396 (2014).

40. N. Romberg et al., Mutation of NLRC4 causes a syndrome of enterocolitis and autoinflammation. Nature genetics 46, 1135–1139 (2014).

41. R. E. Vance, The naip/nlrc4 inflammasomes. Current opinion in immunology 32, 84–89 (2015).

42. C. Duggan et al., Dynamic localization of a helper NLR at the plant–pathogen interface underpins pathogen recognition. Proceedings of the National Academy of Sciences 118, e2104997118 (2021).

43. Y. Zhang et al., A highly efficient agrobacterium-mediated method for transient gene expression and functional studies in multiple plant species. Plant communications 1, (2020).

44. R. Martin, F. Liu, B. Staskawicz, Isolation of Protein Complexes from Tobacco Leaves by a Two-Step Tandem Affinity Purification. Current Protocols 2, e572 (2022).

45. A. Punjani, J. L. Rubinstein, D. J. Fleet, M. A. Brubaker, cryoSPARC: algorithms for rapid unsupervised cryo-EM structure determination. Nature methods 14, 290–296 (2017).

46. E. F. Pettersen et al., UCSF ChimeraX: Structure visualization for researchers, educators, and developers. Protein Science 30, 70–82 (2021).

47. T. D. Goddard et al., UCSF ChimeraX: Meeting modern challenges in visualization and analysis. Protein Science 27, 14–25 (2018).

48. E. C. Meng et al., UCSF ChimeraX: Tools for Structure Building and Analysis. Protein Science, e4792 (2023).

49. P. Emsley, K. Cowtan, Coot: model-building tools for molecular graphics. Acta crystallographica section D: biological crystallography 60, 2126–2132 (2004).

