## Supplementary Materials for "The activated plant NRC4 immune receptor forms a hexameric resistosome"

### The plant "helper" immune receptor NRC4 forms a hexameric resistosome to initiate immune response

**This PDF file includes:**

Materials and methods
Supplemental Text
Figs. S1 to S8
Table S1
References

#### Materials and Methods

Plant materials and growth conditions

Wild-type *N. benthamiana* plants were cultivated in a controlled growth chamber, maintaining an 8-hr-light/16-hr-dark photoperiod at 23°C. For HR assay, 4-week-old *N. benthamiana* plants were used. The *Agrobacterium* strains containing the plasmids were infiltrated into leaves from the abaxial side, and HR phenotype photos were observed and photographed 2 days post-infiltration.

Bacterial strains and growth conditions

The binary constructs were transformed into *Agrobacterium tumefaciens* strain AGL1, followed by cultivation in YEB (Yeast Extract Broth) at 28 °C. Antibiotics were added to the growth medium at the following concentrations (in mg/L): Kanamycin 50, Rifampicin 100, and carbenicillin 100.

Transient expression and western blotting in *N. benthamiana*

Proteins were expressed in leaves of 4-5-week-old *N. benthamiana* plants through transient agroinfiltration. The agrobacterium culture were prepared as described (*1*). For Western blot analysis, two 10-mm leaf discs were collected for each sample 2 days after infiltration. The samples were immediately frozen in liquid nitrogen and ground into a fine powder. The ground samples were then resuspended in 200 µl of Piece™ IP-lysis buffer (Thermo Fisher Scientific, 87787) with 1X protease inhibitor cocktail (Thermo Fisher Scientific, 1861279). With gentle rotation for 15 mins at 4°C, total proteins were denatured by supplement with proper volume of 4X Laemmli Sample buffer (Bio-Rad, 1610747). After centrifugation at 16000g for 15 mins, the supernatants were resolved using SDS-PAGE followed by immunoblotting analysis using HRP-conjugated anti-StrepTag™II antibody (Sigma, 71591-M). In the co-immunoprecipitation assay, Strep-Tactin® XT 4Flow® high-capacity resin was used for the protein enrichment. The resins with immunoprecipitated proteins were then suspended in proper volume of 4X Laemmli Sample buffer with heating at 95°C for 10 mins to elute proteins. Eluted proteins were subsequently resolved using SDS-PAGE, followed by immunoblotting analysis using HRP-conjugated ANTI-FLAG M2 monoclonal antibody (Sigma, A8592). Photography of HRP signal was carried out using ProSignal® Femto ECL reagent (Genesee Scientific). InstantBlue® (abcam) Coomassie-stained membranes were photographed as loading controls for a similar amount of total proteins.

Protein expression and purification

Protein purification was conducted following a modified protocol based on a previously published method (*2*). Vectors containing the NRC4 L9E/DV, genetically fused to either a C-terminal 3Xflag or twin-Strepii, were transformed into *Agrobacterium* AGL1. They were transiently co-expressed in the presence of the P19 suppressor in 4-5-weeks-old *N. benthamiana* leaves by agroinfiltration. Leaf tissue was harvested 2 days after infiltration, ground using a mortar and pestle, and then resuspended in a buffer containing supplemented with protease inhibitors as described (*2*). The lysate was further processed by sonication. The soluble fraction of the lysate was separated from cell debris by centrifugation at 20000g for 45 min at 4°C. The supernatant was loaded onto a column with Macro-Prep DEAE Resin (Bio-Rad Laboratories) using a peristaltic pump (Gilson Incorporated). Sequential affinity chromatography was performed, first with Flag-immunoprecipitation (ANTI-FLAG M2 Affinity Gel, Millipore Sigma), and then Strep-Tactin immunoprecipitation (Strep-Tactin® XT 4Flow® high capacity resin, IBA), to capture the protein complex. The purified proteins were subsequently analyzed using SDS-polyacrylamide gel electrophoresis (PAGE), as shown in Fig.S2.

Cryo-EM sample preparation and data collection

For Cryo-EM sample preparation, Quantifoil Au 2/1 grids with a 2nm carbon layer or Quantifoil Au 1.2/1.3 grids coated with a graphene layer were utilized. The grids were glow discharged twice (25 + 10 seconds) at 15 mA. Subsequently, 3µl of freshly prepared NRC4 L9E/DV protein was applied to the grids, followed by blotting for 5 seconds at 4 °C under 100% humidity conditions. The grids were then rapidly plunge-frozen in liquid ethane using an FEI Vitrobot Marked IV (Thermo Fisher Scientific). Data collection was carried out using a Titan Krios electron microscope, operated at 300 kV and equipped with a K3 direct electron detector camera situated behind a BioQuantum energy filter. A total of 13,370 (carbon layer) and 14,400 (graphene layer) dose-fractionated movies were collected, with an electron exposure of 50 electrons per Å$^2$ and a defocus range of -1.0 µm to -2.0 µm. All movies were acquired in super-resolution mode, with a pixel size of 0.525 Å.

Cryo-EM data processing

Cryo-EM data for the NRC4 resistosome was processed using CryoSPARC (*3*). Initial steps included Patch Motion Correction and Patch CTF Estimation (*3*).

Particles were picked using Blob Picker, extracted into 100-pixel boxes, after binned by a factor of 4 to accelerate data processing. Following several rounds of 2D classification, clear top views of the NRC4 complex and two distinct side views were identified, making apparent the presence of both hexameric and dodecameric NRC4 resistosome assemblies. To specifically pick particles for the hexameric or the dodecameric NRC4 resistosomes, two corresponding templates were selected and used in a round of template-bade picking. Particles representing the hexameric resistosome were extracted using a 90-pixel box with a pixel size of 4.2 Å. After several rounds of 2D classification, classes displaying clear secondary structural features were selected. This yielded a total of 166,926 particles, which were re-extracted into 360-pixel boxes with a pixel size of 1.05 Å. An initial model was generated through Ab-initio Reconstruction with C1 symmetry. After two rounds of Heterogeneous Refinement with C1 symmetry, 80,368 particles were chosen for Non-uniform Refinement with C6 symmetry. This procedure resulted in a cryo-EM map at 2.81 Å resolution. The cryo-EM map was further improved through Defocus Refinement and Global CTF Refinement, resulting in a final resolution of 2.66 Å that was used for model building (Fig.S3, S7 and S8).

Particles representing the hexameric resistosome were extracted using 100-pixel boxes after binning by 4 to accelerate data processing. Multiple rounds of 2D classification were employed to remove damaged particles or contaminants, while retaining as many "good" particles as possible, even if some exhibited lowered resolution features. 1,068,393 particles were subsequently re-extracted using a 400-pixel box and a pixel size of 1.05 Å. An initial model was constructed through Ab-Initio Reconstruction with C1 symmetry, providing an initial, low-resolution view of the double-layer resistosome. The dataset was partitioned into two subsets, each subjected to three rounds of Heterogeneous Refinement with C1 symmetry. The best classes resulting from this process were chosen and further cleaned through 2D classification. The "best" particles (28,066) displaying clear secondary structural features were selected and refined to a resolution of 3.64 Å through Non-uniform Refinement with D6 symmetry. The resolution was further improved by Defocus Refinement and Global CTF Refinement, resulting in a 3.54 Å cryo-EM map that was used for subsequent model building and structural interpretation (Fig.S3 and S7).

Model building and validation

The structural models of the five NRC4 protein domains, namely CC, NBD, HD1, WHD and LRR, were initially predicted using AlphaFold (*4, 5*). Subsequently, each domain structure was docked into the hexameric NRC4 cryo-EM map using rigid body fitting in ChimeraX (*6-8*). Atomic models were manually adjusted residue by residue in COOT (*9*). A similar approach was followed for the dodecameric NRC4 model assembly. In this case, AlphaFold multimer was used to predict the dimerized structure the $\alpha$1-helix of the CC domain, which was then docked into the cryo-EM map through rigid body fitting. The resulting models were refined against the hexameric and dodecameric NRC4 cryo-EM density maps using real space refinement in PHENIX, respectively. The statistics of cryo-EM data collection, processing, model building, and validation are listed in Table.S1. Figures were generated using ChimeraX (*6-8*) and pyMOL (Schrödinger).

<u>In planta $Ca^{2+}$ influx assay</u>

*Agrobacterium* strains were syringe-infiltrated into *N. benthamiana* leaves expressing GCaMP3. After a 5-hour incubation, leaf discs with a 0.4 cm diameter were carefully collected using a cork borer and transferred to a 96-well flat bottom black Corning assay plate containing 200 μL distilled water in each well. The plate was then kept on bench for 1 h to reset $[Ca^{2+}]_{cyt}$ evoked by the sampling process, followed by fluorescence recording of GCaMP3 at 45 second intervals by a Perkin-Elmer envision multilabel plate reader. GCaMP3 emission was collected using an FITC top mirror at 535 nM with excitation light at 485 nM. The data plotted in the graphs represent the average values obtained from replicate leaf discs. For external $Ca^{2+}$ treatment, collected leaf discs were transferred to the plate containing either distilled water or 10 mM $CaCl_2$ solution, followed by 1 h equilibration and $Ca^{2+}$ recoding procedure as describe above. For $Ca^{2+}$ blocker treatment, Agrobacteria with 2 mM $LaCl_3$ in infiltration buffer were co-infiltrated into *N. benthamiana* leaves for the subsequent $Ca^{2+}$ recording procedure.

Cell viability assays

HEK293FT cells were seeded at $4\times10^5$ cells/well in 12-well plates for transfection. The following day, cells were transfected with plasmids encoding mCherry, NRG1.1$^{DV}$-mCherry, and NRC4$^{DV}$-mCherry using Mirus TransLT1 Transfection Reagent (Mirus, MIR2300) following the manufacturer's instructions. Individual wells were imaged over time at 10x magnification with an Incucyte System (Sartorius) in a 37°C, 5% $CO_2$ incubator. Protein expression was tracked by mCherry fluorescence, and cell confluence per image was calculated using Incucyte Analysis Software.

HEK293FT cells were plated at $2\times10^4$ cells/well in 96-well plates and transfected with mCherry, NRG1.1$^{DV}$-mCherry, and NRC4$^{DV}$-mCherry plasmids using Mirus TransLT1 Transfection Reagent (Mirus, MIR2300). Cell viability was measured at 24 h and 48 h post transfection using CellTiter-Glo (Promega) according to the manufacturer's instructions.

Fluorescent microscopy assay

HEK293FT cells were seeded on 8-well chamber slides and transfected with mCherry, NRG1.1$^{DV}$-mCherry, and NRC4$^{DV}$-mCherry plasmids using Mirus TransLT1 Transfection Reagent (Mirus, MIR2300). Cells were fixed with 4% PFA at 48 h post transfection and stained with Wheat Germ Agglutinin (Alexa Fluor 488, Invitrogen) and Hoechst for 10 min at room temperature. Subsequently, cells were washed with PBS and mounted using Vectashield mounting medium (Vector Laboratories, H-1000). Images were acquired with an inverted confocal microscope (Zeiss LSM 800) and processed with ZEN software (Zeiss). Image analysis was performed with the ImageJ software.

#### References and notes


1. Y. Zhang *et al.*, A highly efficient agrobacterium-mediated method for transient gene expression and functional studies in multiple plant species. *Plant communications* **1**, (2020).
2. R. Martin, F. Liu, B. Staskawicz, Isolation of Protein Complexes from Tobacco Leaves by a Two-Step Tandem Affinity Purification. *Current Protocols* **2**, e572 (2022).
3. A. Punjani, J. L. Rubinstein, D. J. Fleet, M. A. Brubaker, cryoSPARC: algorithms for rapid unsupervised cryo-EM structure determination. *Nature methods* **14**, 290-296 (2017).
4. M. Varadi *et al.*, AlphaFold Protein Structure Database: massively expanding the structural coverage of protein-sequence space with high-accuracy models. *Nucleic acids research* **50**, D439-D444 (2022).
5. J. Jumper *et al.*, Highly accurate protein structure prediction with AlphaFold. *Nature* **596**, 583-589 (2021).
6. E. F. Pettersen *et al.*, UCSF ChimeraX: Structure visualization for researchers, educators, and developers. *Protein Science* **30**, 70-82 (2021).
7. T. D. Goddard *et al.*, UCSF ChimeraX: Meeting modern challenges in visualization and analysis. *Protein Science* **27**, 14-25 (2018).
8. E. C. Meng *et al.*, UCSF ChimeraX: Tools for Structure Building and Analysis. *Protein Science*, e4792 (2023).
9. P. Emsley, K. Cowtan, Coot: model-building tools for molecular graphics. *Acta crystallographica section D: biological crystallography* **60**, 2126-2132 (2004).

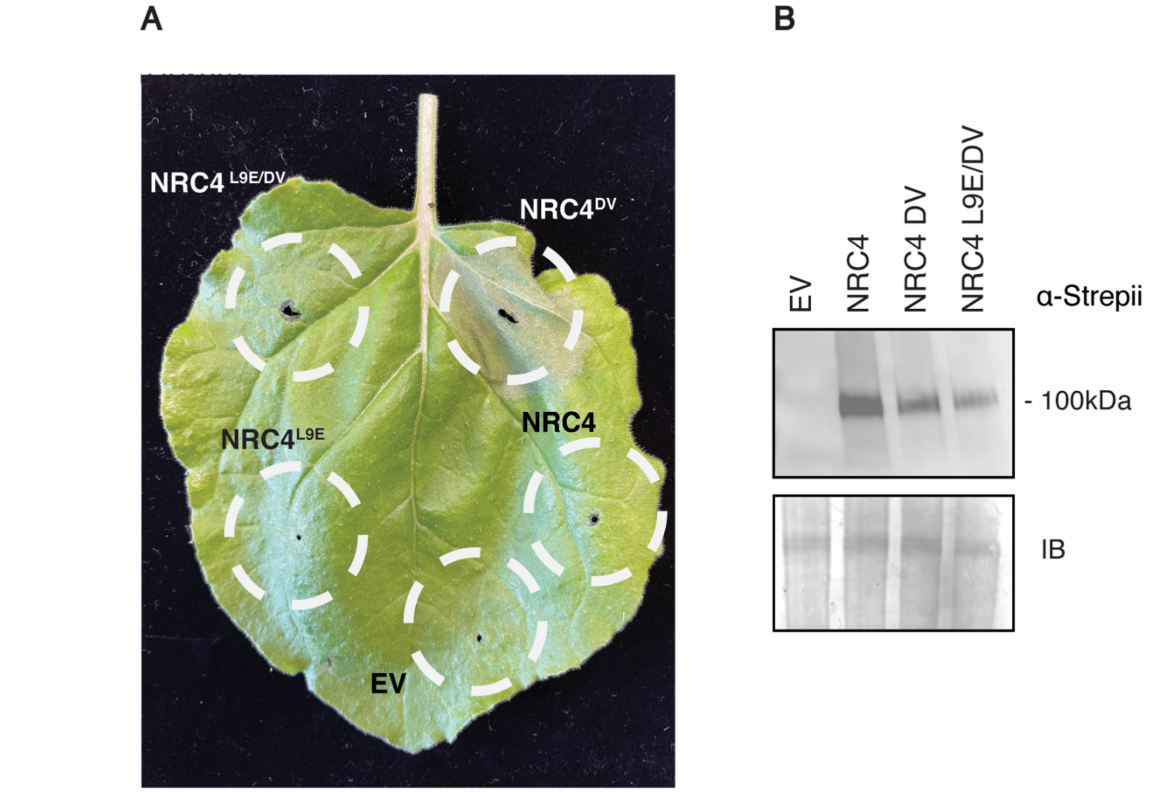


**Fig.S1. Auto-active NRC4 DV triggers cell death and NRC4 L9E/DV impairs cell death. (A)** *N. benthamiana* leaves infiltrated with *Agrobacterium* harboring the indicated Strepii-tagged constructed to observe cell death phenotype 2 days post infiltration. **(B)** Strepii-tagged proteins extracted from harvested leaf tissue 2 days post infiltration were resolved by SDS-PAGE and blotted for α-Strepii. InstantBlue® Coomassie-stained gel as loading control.

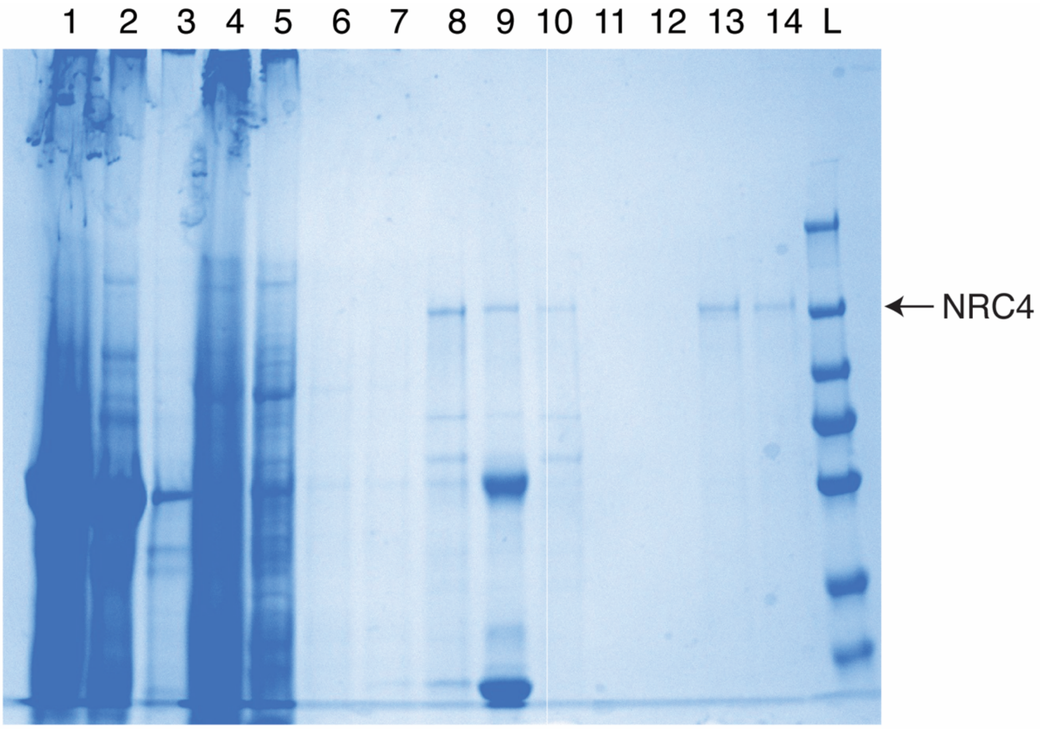


1: Lysed cells
2: Flowthrough after binding DEAE resin (Macro-Prep DEAE Resin, Bio-Rad)
3: Wash
4: Elution from DEAE resin
5: Flowthrough after binding to ANTI-FLAG M2 affinity gel (Sigma-Aldrich)
6-7: Successive washes of ANTI-FLAG M2 affinity gel
8-9: Sucessive elution from ANTI-FLAG M2 affinity gel
10: Flowthrough after binding to Strep-Tactin XT 4Flow high capacity resin (iba)
11-12: Successive washes of Strep-Tactin XT 4Flow high capacity resin
13-14: Successive elutions of Strep-Tactin XT 4Flow high capacity resin
L: Ladder (PageRuler Prestained Protein Ladder, Thermo Finsher)

**Fig.S2. Purification of the NRC4 L9E/DV complex.** Samples were analyzed by SDS-PAGE and Instantblue® Coomassie staining.

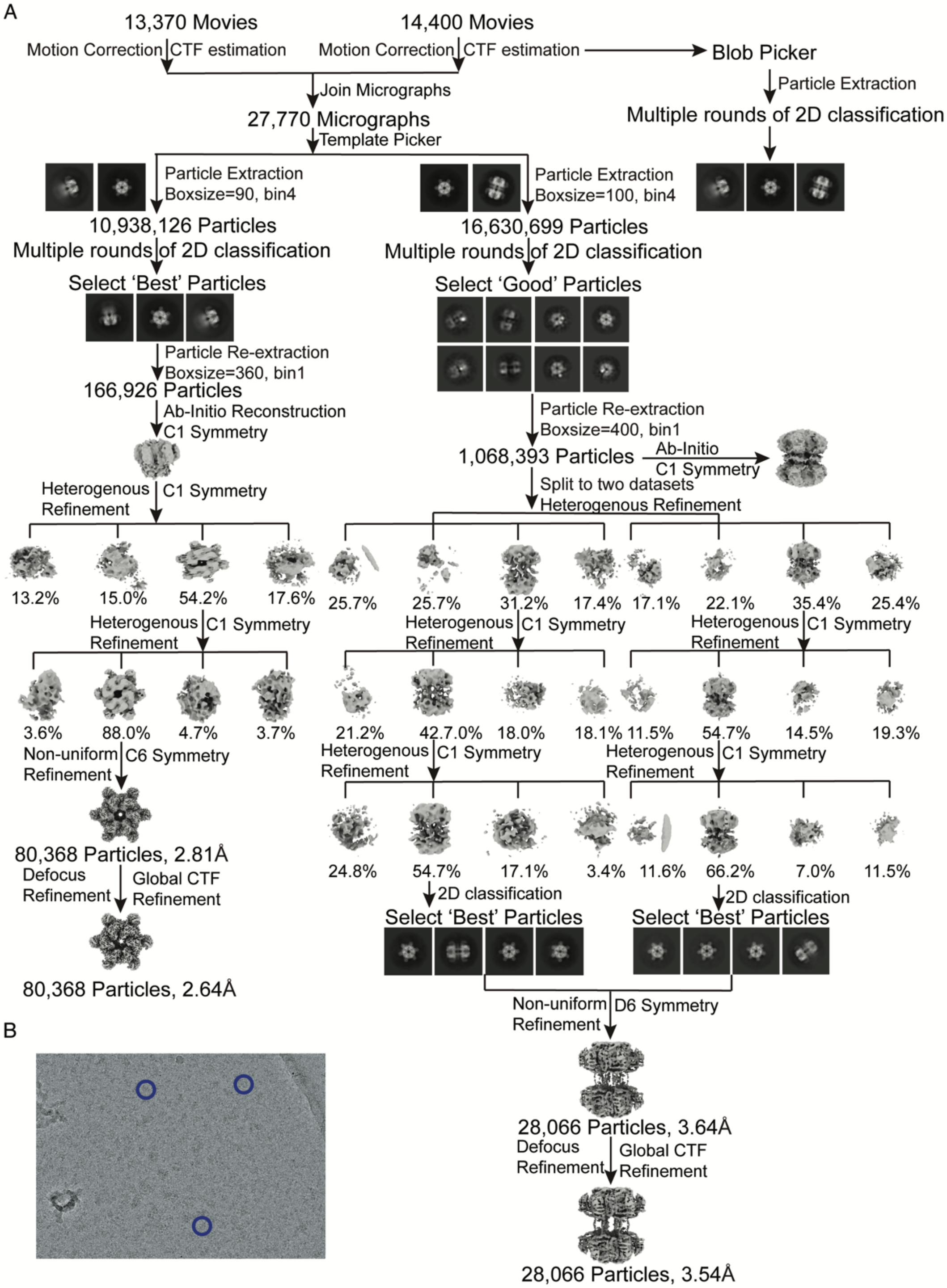


**Fig.S3. Cryo-EM data processing workflow. (A)** Data processing tree from collected movies to the reconstruction of the NRC4 resistosome. **(B)** A representative cryo-EM image of the NRC4 dataset. NRC4 particles are indicated by blue circles.

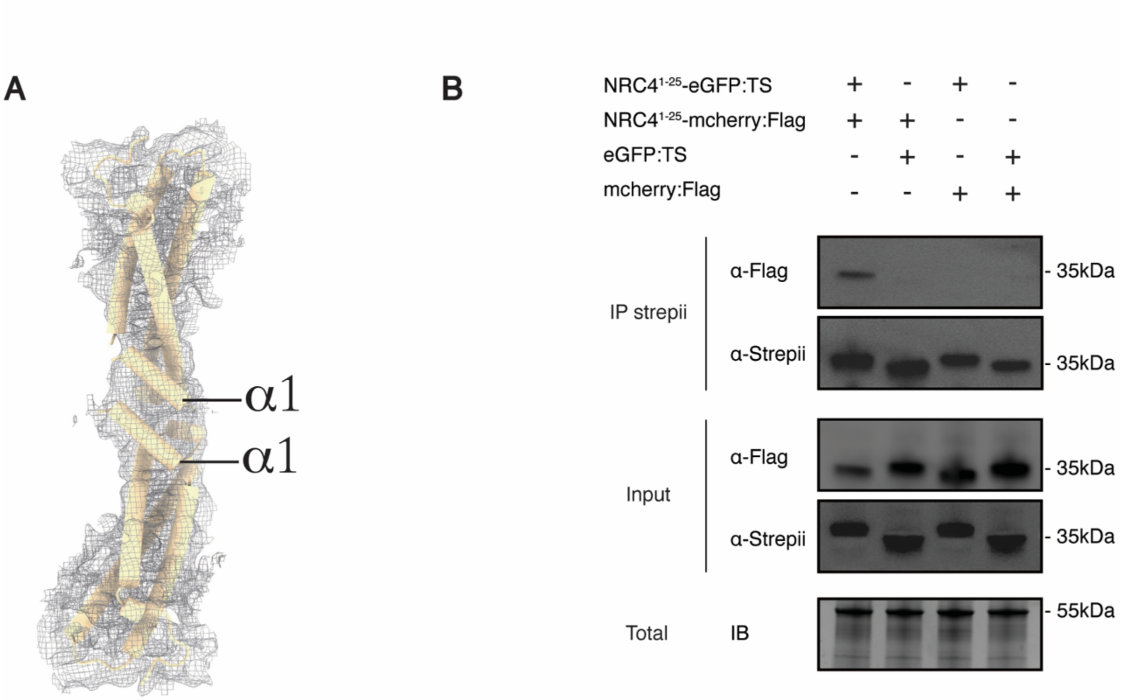


**Fig.S4. Structural detail of the dual-hexamer interface. (A)** Cryo-EM map of the two CC domains originating from adjacent protomers, situated on opposite sides of the double-layer hexamer. The CC domains play a crucial role in mediating the formation of the dodecameric NRC4 resistosome, where the α1-helices interact with their counterparts from the opposite side. **(B)** The co-IP assay confirming the interaction between the two α1-helices (1-25). Agrobacterium stains harboring designated vectors were co-infiltrated into *N. benthamiana* leaves. Subsequent co-IP experiments utilized Strep-Tactin® XT 4Flow® high-capacity resin for protein extraction, followed by analysis through protein gel blotting with α-Flag or α-Strepii antibodies. InstantBlue® Coomassie-stained gel as loading control (IB, InstantBlue; IP, immunoprecipitation.)

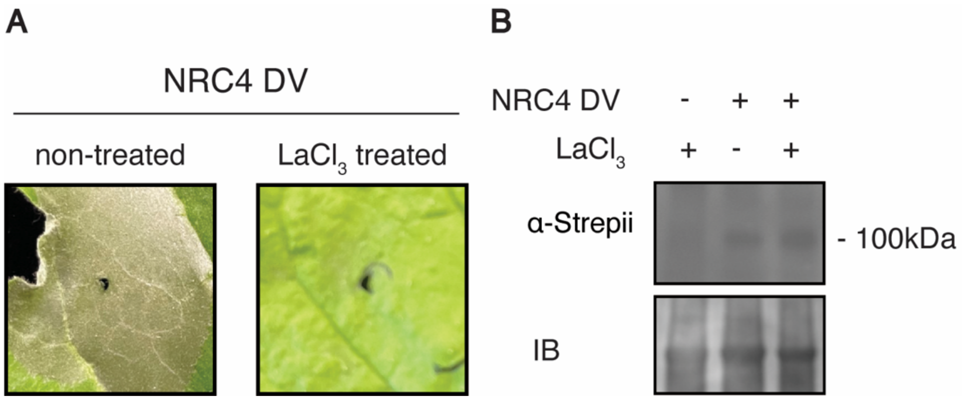


**Fig. S5. (A)** Hypersensitive response phenotypes of *N. benthamiana* leaves upon the expression of active NRC4 DV in the absence and presence of the $Ca^{2+}$ channel blocker $LaCl_3$ (2mM). **(B)** The protein expression levels of NRC4 DV in the *N.benthamiana* leaves were evaluated using SDS-PAGE and subsequent immunoblotting with α-Strepii antibody. InstantBlue® Coomassie-stained gel as loading control.

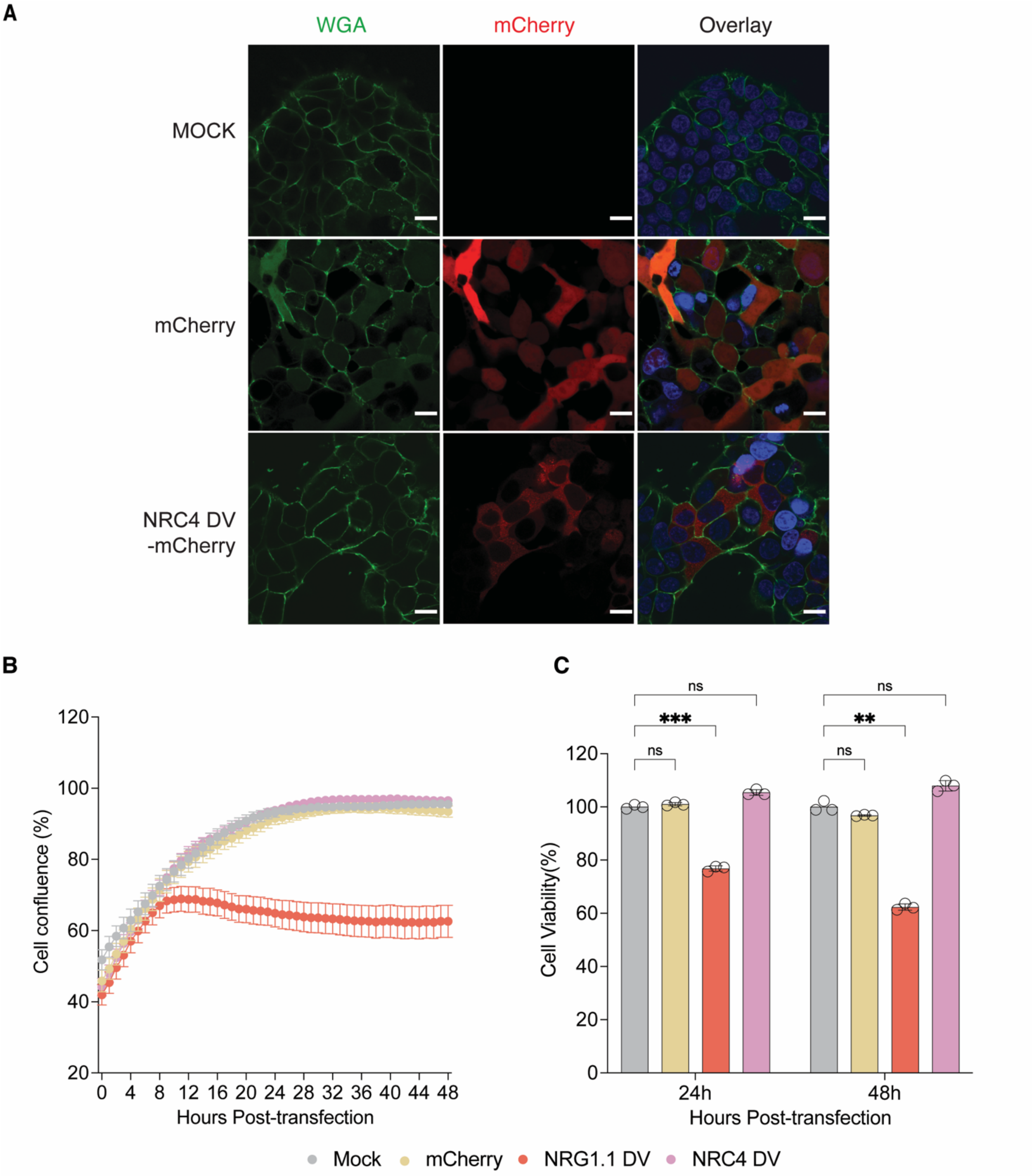


Fig.S6 **(A)** Confocal imaging of mock, mcherry, and NRC4 DV-mCherry transfected 293FT cells at 48 h post transfection. Red, mCherry. Green, Wheat germ agglutinin (WGA). Blue, hoeches nuclear stain. Scale bars, 20 µm. **(B)** Time course of cell confluence for mock, mCherry, NRG1.1 DV-mCherry, and NRC4 DV-mCherry transfected 293FT cells (mean ± SEM, n=9). **(C)** Cell viability of mock, mCherry, NRG1.1 DV-mCherry, and NRC4 DV-mCherry transfected 293FT cells at 24 h and 48 h post transfection using the CellTiter-Glo luminescent cell viability assay (mean ± SEM, n=3).

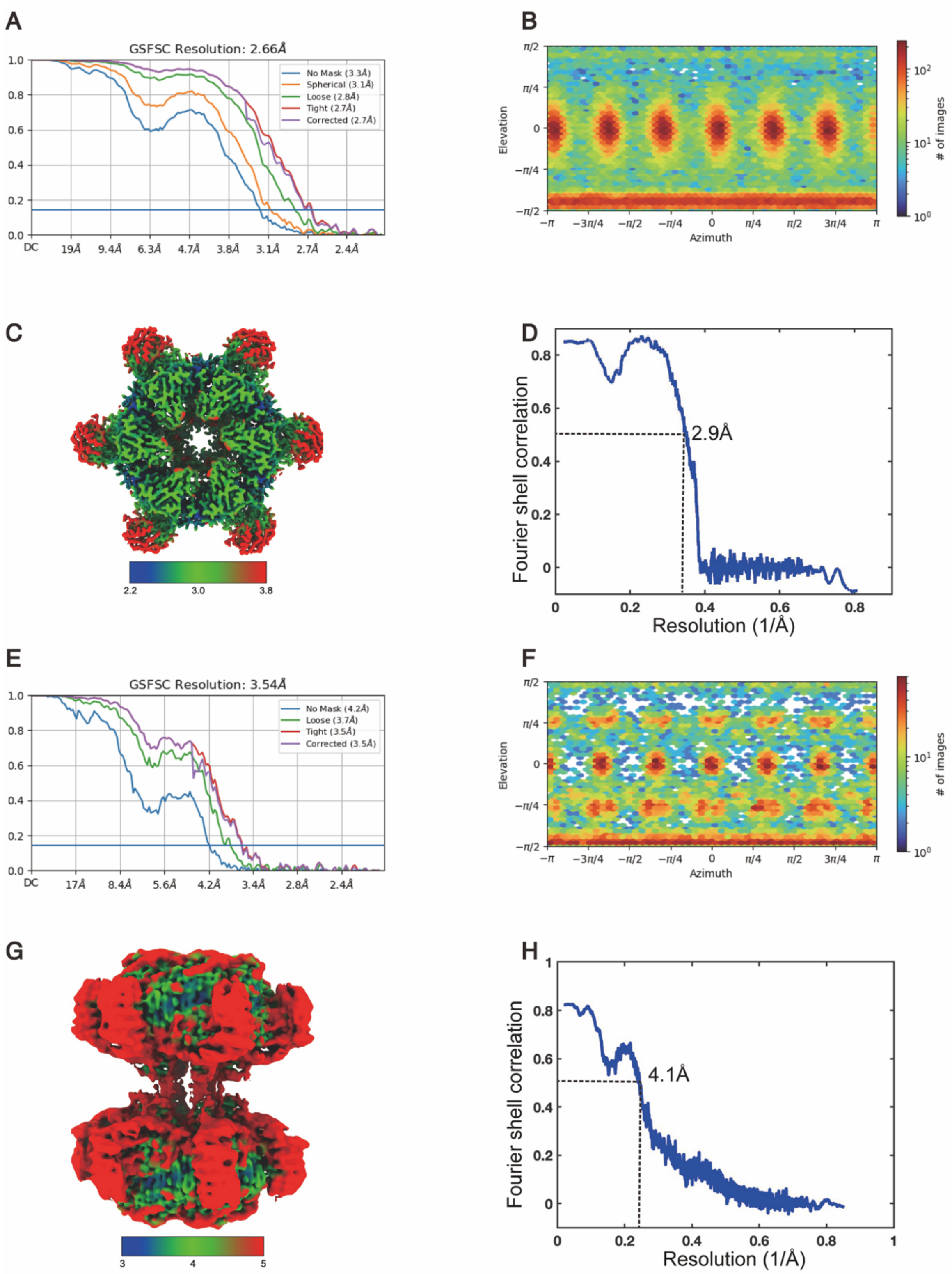


**Fig.S7. Map quality of the NRC4 resistosome. (A-D)** GSFSC curve, angular distribution of the final 3D reconstruction, local resolution and model vs. map FSC plot, respectively, for the NRC4 hexamer. **(E-H)** GSFSC curve, angular distribution of the final 3D reconstruction, local resolution and model vs. map FSC plot, respectively, for the NRC4 dodecamer.

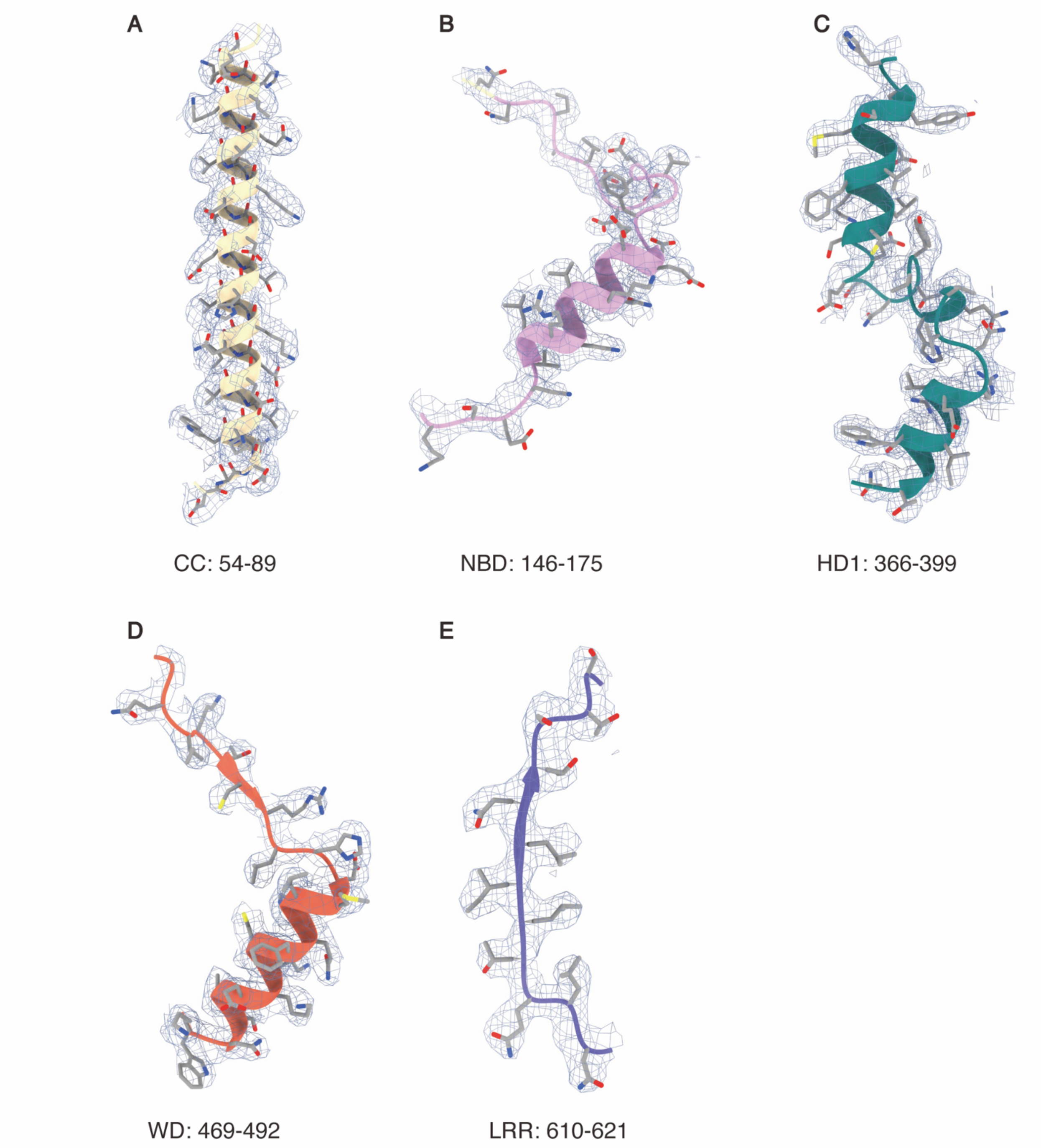


**Fig. S8.** Local cryo-EM density maps of representative regions of the NRC4 resistosome shown with fitted atomic models

**Table S1. Cryo-EM data collection, processing and refinement.**

| **Data collection and processing** | NRC4 Hexamer (PDB:XXX EMDB:XXX ) | NRC4 Dodecamer (PDB:XXX EMDB:XXX ) |
|---|---|---|
| Microscope | Krios | Krios |
| Camera | K3 | K3 |
| Voltage (kV) | 300 | 300 |
| Camera Magnification | 81,000 | 81,000 |
| Pixel size (Å) | 1.05 | 1.05 |
| Electron exposure (e-/Å2) | 50 | 50 |
| Number of frames per exposure | 50 | 50 |
| Defocus range (µm) | -1 ~ -2 | -1 ~ -2 |
| Automation software | SerialEM | SerialEM |
| Micrographs collected (no.) | 27,770 | 27,770 |
| Refined particles (no.) | 80,368 | 28,066 |
| Map resolution (Å) | 2.64 | 3.54 |
| Map resolution range (Å) | 2.2 - 3.8 | 3.0 - 5.0 |
| Sharpening tool | cryoSPARC | cryoSPARC |
| Map sharpening B factor (Å2) | 88.7 | 95.4 |
| **Model Composition** | | |
| Ligand | 6 | 12 |
| Protein | 6 | 12 |
| Residues | 5028 | 10194 |
| **Refinement** | | |
| Refinement package | Phenix | Phenix |
| Real or reciprocal space | Real | Real |
| Resolution cutoff (Å) | 2.64 | 3.54 |
| Model-Map scores | | |
| CC | 0.74 | 0.65 |
| FSC 0.5 (Å) | 2.9 | 4.1 |
| B factors (Å2) | 67.99 | 117.85 |
| R.M.S deviations | | |
| Bond length (Å) | 0.005 | 0.006 |
| Bond angles ($^{o}$) | 0.648 | 0.760 |
| Validation | | |
| MolProbity score | 1.99 | 2.2 |
| CaBLAM outliers | 1.78 | 2.61 |
| Clash score | 6.94 | 19.67 |
| Ramachandran plot | | |
| Favoured (%) | 95.49 | 93.82 |
| Allowed (%) | 4.23 | 5.92 |
| Outliers (%) | 0.28 | 0.26 |

Table S1. Statistics of the cryo-EM data collection, processing and refinement.
